# Epiphenomenal neural activity in the primate cortex

**DOI:** 10.1101/2022.09.12.506984

**Authors:** Sébastien Tremblay, Camille Testard, Jeanne Inchauspé, Michael Petrides

## Abstract

When neuroscientists record neural activity from the brain, they often conclude that neural responses observed during task performance are indicative of the functional role of the brain area(s) studied. In humans and nonhuman primates, it is often hard to combine recordings and causal techniques within the same experiment, leaving the possibility that the activity recorded may be epiphenomenal rather than reflecting a specific functional role. Currently, the prevalence of epiphenomenal neural activity in the cortex is unknown. To estimate the extent of such activity in primates, we chronically recorded neural activity in the prefrontal cortex of the same monkeys using the same neural implants during the performance of four different cognitive tasks. The four tasks were carefully selected such that only one of them causally depends on the brain area recorded, as demonstrated by previous double dissociation studies. Using the four most common single neuron analyses methods in the field, we found that the prevalence and strength of neural correlates were just as high across all four tasks, including for the three tasks that do not depend on this brain area. These results suggest that the probability of observing epiphenomenal activity in primate cortex is high, which can mislead investigators relying on neural recording or imaging to map brain function.

**One-Sentence Summary:** Tremblay, Testard and colleagues show that inferring a brain area’s function from neural recordings alone could be misleading.

---

Recording neural activity from awake behaving animals is a common approach to examine the role of different brain areas in cognition^1–3^. For example, neural responses correlating with decision variables during a task may indicate that the recorded neurons play a functional causal role in those decisions^4^. Using this inferential logic, various roles have been assigned to several areas of the cerebral cortex of mammals, including humans^5–7^. This method, however, only provides *correlational* evidence of the involvement of a brain area in a cognitive process^8^. Although it is now standard in rodents and smaller animals to include causal techniques as supporting evidence in the same study, it is most often impractical to do so in studies involving human and nonhuman primate subjects. Thus, it would be helpful to know how reliable neural correlates uncovered in the primate brain are for estimating the functional role(s) of a brain area.

The inferential power of the brain recording and imaging methods is limited by the prevalence of epiphenomenal neural activity in the brain, that is, neural activity correlated with task variables that plays no necessary role in the performance of that task^9^. One example of epiphenomenal activity is the presence of motor signals in the primary visual cortex^10^. The presence of such epiphenomenal neural responses can mislead investigators when hypothesizing about the functional role of a brain area in health and disease^11,12^. Models based on such evidence can lead to contradictory theories of brain area function^13,14^ and can identify ineffective therapeutic targets in clinical research. Moreover, the number of proposed roles for each brain area inflates over time as more of these neural correlates are uncovered^15^, undermining brain mapping efforts.

In the present study, we sought to investigate the prevalence of epiphenomenal signals in one actively studied part of the primate brain: the prefrontal cortex (PFC). We focused on a sub-region within the PFC, pre-arcuate area 8A, which has a well-defined role based on causal studies in human patients and double dissociation lesion studies in non-human primates^16^. Area 8A of the primate PFC has been demonstrated to be necessary for the conditional selection of visual stimuli based on instruction cues (aka. “top-down” selection), as shown by a severe and permanent deficit following bilateral lesions to this area^17–20^, and a series of microstimulation and inactivation studies^21–24^. The same area plays no causal role in basic visual discrimination, motor control or working memory, as shown by unaltered behavioral performance on appropriate tasks following complete, bilateral 8A lesions^18,19,25,26^. The non-involvement of area 8A in working memory is further supported by the fact that impairment in working memory is observed after inactivation of the immediately adjacent prefrontal peri-principalis area 46, providing a double dissociation of function^18,27–30^. Double dissociations rule out alternative explanations for the observed deficits, such as compensation, plasticity, or task difficulty, and is the gold standard for causal inference in neuropsychology^31^.

Using chronic Utah array neural implants, we recorded simultaneously from large neural populations in area 8A in the same monkeys and using the same implants during the performance of four different cognitive tasks (**Figure 1A, B**). Critically, based on the causal studies mentioned above, only one of the tasks selected depends on area 8A function. The other three tasks were selected because *they do not* depend on area 8A, i.e. performance is not impaired after area 8A lesions (**Figure 1C**). This approach allowed us to compare neural activity during tasks that depend and tasks that do not depend on the recorded brain area. We compared single neuron spiking activity across the four tasks using four standard metrics widely used in the field to uncover the neural basis of cognition: single neuron selectivity^32^, persistent firing activity^33^, signal detection theory (auROC^34^), and neural population decoding^35^. We tested whether differences in neural responses between tasks would reliably reflect the known functional role of this brain area, or if epiphenomenal neural activity would be observable in some of the control tasks.

**Figure 1.**
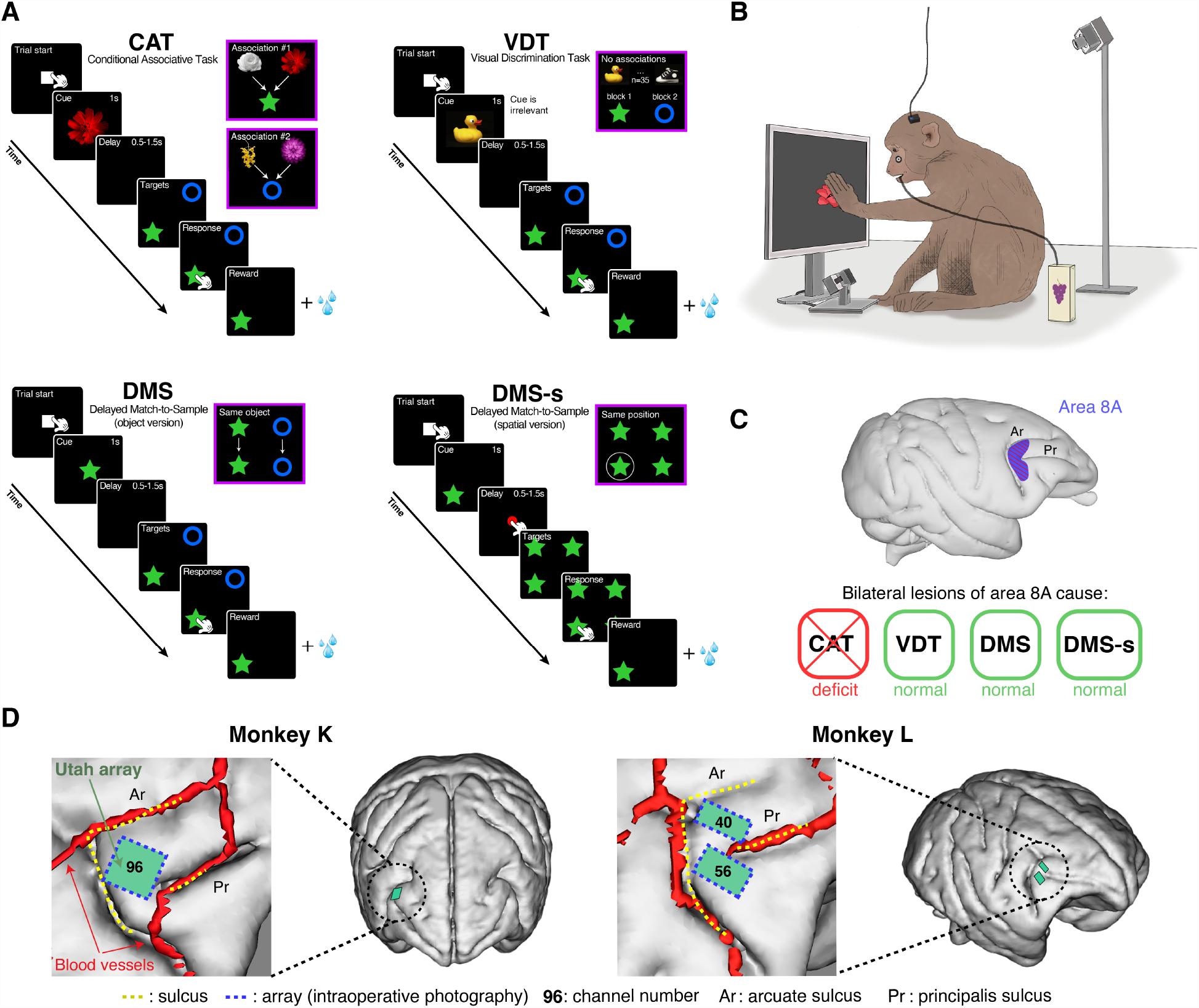
Behavioral tasks, double dissociation of function, and neural recordings. (**A**) Monkeys performed four different cognitive tasks on a touchscreen while neural activity was recorded through the same chronic implants in area 8A. Each task had the same format and included a cue, a delay, and a response epoch. In all tasks, the monkey had to make a selection between a blue circle and a green star target. Thus, the only difference was the cognitive operation to be performed on those two visual target stimuli. The CAT task required selecting the correct visual target based on a previously established conditional relation with the visual instruction cues. Thus, the CAT task measured the instructed selection of a particular visual target, i.e. the instructed allocation of attention to a visual target. The VDT task required selecting the same visual target on every trial, regardless of the visual cue presented, based on a block design, and thus measured the visual perceptual identification of the visual target stimuli. The DMS task required selecting the target that was identical to the visual stimulus presented a short time earlier and thus measured visual short-term memory. The DMS-s task required selecting the target based on the spatial position it occupied a short time earlier and thus measured visual spatial short-term memory. In DMS-s, the delay was initiated after pushing a red circle at the center of the screen to prevent the monkeys from leaving their hands at the cued position. (**B**) Depiction of the experimental setup. Monkeys were not head-restrained and body and eye movements were tracked through video cameras, as in^*38*^. (**C**) Results from lesion studies showing that only CAT performance is severely and permanently affected by bilateral inactivation of area 8A. Performance of VDT, DMS, and DMS-s is not affected. (**D**) Position of chronic Utah array implants within area 8A of both monkeys. Brain and vasculature reconstructed from MRI.

Two macaque monkeys were trained on the following four tasks before the beginning of neural recording sessions: a conditional associative task (CAT), a visual discrimination task (VDT), a delayed match-to-sample task (DMS), and a spatial version of the delayed match-to-sample task (DMS-s). As explained earlier, only the CAT task depends critically on cortical area 8A; the other three tasks involving visual object identification or working memory do not (**Figure 1C**). Behavioral performance of the monkeys was high across all four tasks (>80%) and comparable across the four tasks (**Figure 2A**). We implanted custom-designed Utah arrays that fitted optimally within area 8A based on individual anatomy and vasculature reconstructed from MRI (**Figure 1D**). We recorded simultaneously from an average of 166 single and multi-units in monkey K and 134 in monkey L in each session (40 sessions, 5 per monkey per task, total of 3335 units and 2688 units for monkeys K and L, respectively). The number of recorded neural units was comparable across tasks for each monkey (**Figure 2B**). Only neural activity from the same neural implants was compared across tasks.

**Figure 2.**
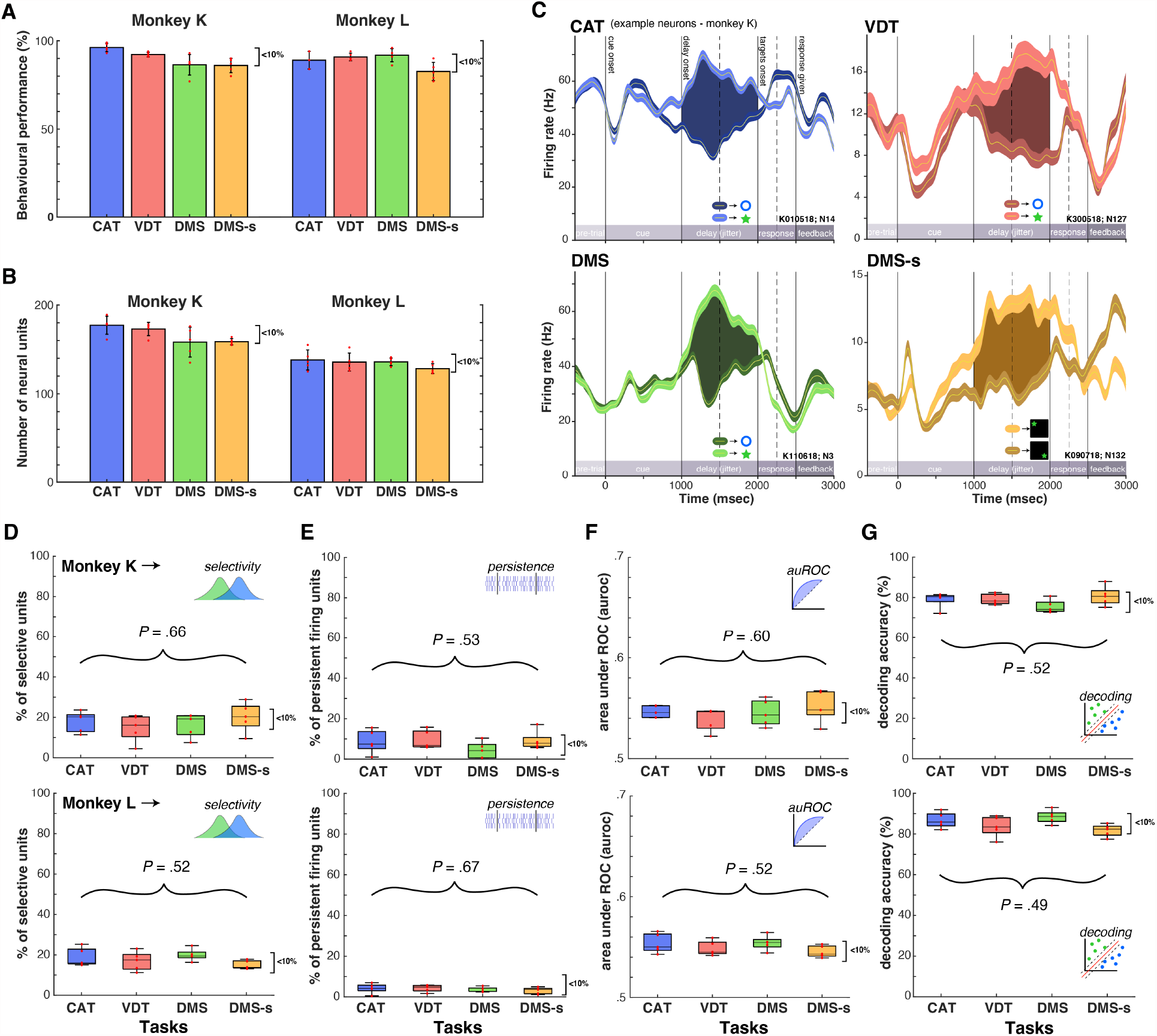
Comparisons of behavioral performance and standard neural activity metrics across tasks. (**A**) Behavioral performance of both monkeys on each of the four tasks. Average performance was comparable across tasks (all medians fall within a 10% interval). Red dots indicate individual sessions. 5 sessions per task. (**B**) Number of neural units (single and multi-units) recorded simultaneously during each task. The average number of units recorded was similar across tasks for each monkey. (**C**) Example single neuron spiking activity during the performance of each task, averaged over trials, for monkey K. Trials are separated based on whether target 1 or target 2 needed to be selected on this trial. This selection depended on the cognitive task. Shaded area shows significant tuning during the delay epoch of each task, when no stimuli are on the screen. Error bars are SEM. (**D**) Proportion of neurons showing significant tuning during the delay, for each task. *P* values represent one-way ANOVA comparing means of the four tasks, corrected for uninstructed movements (see **Methods**). (**E**) Proportion of persistent firing units during the delay epoch, for each task. *P* values as in (D). (**F**) Area under the receiver operating curve (auROC) computed for every unit, averaged across units. *P* values as in (D). (**G**) Single-trial decoding accuracy of a support vector machine predicting target selection during the delay epoch. *P* values as in (D). Box plots in (D-G) represent median and the 25^th^ and 75^th^ percentiles. Each red dot is a session. *P*-values are FDR-corrected.

We focused the analyses on a key epoch that is comparable across tasks: the *delay epoch* (see **Methods**). During that epoch, there are no stimuli on the screen and the monkey cannot prepare a motor response because of the randomization of target locations in the following epoch (except in the case of the DMS-s where motor responses could be planned). For these reasons, in the *delay epoch* the “cognitive” representations are dissociated from direct sensory and/or motor representations. In the cognitive neurophysiology field, functional roles of brain areas are typically inferred from the presence of cognitive neural signals during this delay epoch^36^.

Each of the four tasks used taps into a dissociable cognitive process during the delay epoch: conditional visual selection, i.e. the top-down instructed allocation of attention to a visual stimulus, in CAT, visual perceptual discrimination in VDT, working memory for visual object in DMS, and working memory for visual space DMS-s; see **Methods**). Thus, neurons must encode different cognitive representations to support behavioral performance in the delay epoch. We tested whether neurons in area 8A only encoded the representation required for the CAT task, or whether they also encoded the representations required by the other tasks that do not depend on area 8A. Surprisingly, we found evidence of single neuron selectivity to the cognitive representation required by all four tasks in both monkeys (**Figure 2C & Figure S1**).

Although example single neuron recordings can provide important information, the field typically considers the proportion of all neurons that have similar task selectivity to support hypotheses about function^37^. We thus compared the proportion of tuned neurons during the delay epoch across all tasks relative to the CAT task. After correcting for task-aligned uninstructed movements that inflate selectivity^10,38^, we found no difference in the proportion of tuned neurons across tasks in either monkey (**Figure 2D**, see **Methods**). Note that in Monkey L who did not exhibit uninstructed movements correlated with task variables, results are identical with or without movement correction (**Supp. Fig. 2, 3 & 4**). In all four tasks, 17.3% and 17.8% of neurons recorded during a session exhibited statistically significant selectivity during the delay epoch, for monkey K and L, respectively (one-way ANOVA F(3,16) = 0.68 and F(3,16) = 1.66, *P*(two-sided, FDR corrected) = 0.66 and *P* = 0.52 for monkey K and L, respectively).

Another standard neural activity metric widely thought to represent cognitive processing is persistent firing activity during the delay^39^ (although debated, see^40^). Thus, we tested whether differences would arise between tasks when the temporal persistence of neural selectivity is compared. We used a standard criterion for defining persistent activity: significant selectivity during at least 5 consecutive 100 msec bins during the delay (*P* < 0.01; minimal delay length = 500msec). Example neurons that would satisfy this criterion are displayed in **Figure 2C**. When quantifying the proportion of all neurons satisfying this persistence criterion across the four tasks, we found no significant differences across tasks (**Figure 2E**, ANOVA F(3,16) = 1.23 and F(3,16) = 0.52, *P*(two-sided, FDR corrected) = 0.53 and *P* = 0.67 for monkey K and L, respectively, movement corrected as above). In monkey K and L, 7.9% and 3.7% of neurons exhibited persistent activity during the delay, respectively, regardless of whether the task depends on area 8A (CAT), or not (VDT, DMS, & DMS-s).

The principles of signal detection theory have traditionally been applied to characterize the selectivity of individual neurons^41^. One metric from this framework that is commonly used is the area under the receiver operating curve, or auROC, which allows quantifying the selectivity of a single neuron assuming it was a perfect observer. We computed the auROC for each recorded neuron and computed the average auROC across the sample of neurons recorded for every session. When comparing the mean auROC across tasks, we found that they were comparable, a result replicated across monkeys (**Figure 2F**, ANOVA F(3,16) = 0.94 and F(3,16) = 1.46, *P*(two-sided, FDR corrected) = 0.60 and *P* = 0.52 for monkey K and L, respectively, movement corrected). The mean auROC values for monkey K and L were 0.54 and 0.55, respectively, when including selective and non-selective neurons.

Single neuron selectivity and persistent selectivity metrics are based on average firing rates over trials. Although these metrics are informative, they ignore some important neural dynamics unfolding at the population level on a moment-by-moment basis^42^. As the field is developing an appreciation for those dynamic features, more investigators are focusing on single-trial analyses from simultaneously recorded neural ensembles^43^. A popular approach is to use decoding algorithms, such as support vector machines, to decode representations from ensemble activity on single trials^44,45^. Using this approach, we compared the decoding accuracy for target selection during the delay across the four tasks using the activity of simultaneously recorded neural ensembles. Our analyses, restricted to correct trials, revealed no difference in the single-trial decoding accuracy across tasks (**Figure 2G**, ANOVA F(3,16) = 1.75 and F(3,16) = 2.99, *P*(two-sided, FDR corrected) = 0.52 and *P* = 0.49 for monkey K and L, respectively, movement corrected). All four cognitive representations specific to each task were decoded with an average of 78.6% and 85.13% accuracy from simultaneous neural activity during the delay in monkey K and L, respectively (SVM, cross-validated k-fold = 5, see **Methods**).

Neural recordings are powerful approaches to investigate the neural correlates of cognitive processing. However, the reliability of these correlates for revealing the functional role(s) of brain areas in the primate brain is uncertain. Our results demonstrate that strong neural selectivity was present during the delay epoch both at the single and neural ensemble levels, even in cases where the neural activity is known not to be necessary for task performance. This was true both when selectivity is calculated using standard ‘average-over-trials’ metrics and when using single-trial decoding methods. Importantly, these epiphenomenal cognitive signals were present across *all* of the three control tasks tested. Although more tasks should be tested to better estimate the prevalence of epiphenomenal activity, we can conclude from the current study that the probability of observing credible epiphenomenal signals in the primate cortex is high. This implies a high likelihood that functional roles assigned to particular brain areas are misattributed when experimenters rely solely on recording or imaging results from a single task.

It remains possible that differences in neural selectivity across tasks occur at a different task epoch than the delay or require more sophisticated analytical approaches to be detected. For the purpose of this study, we intentionally limited our analyses to standard methods most commonly used in the field of cognitive neurophysiology in primates and the delay epoch during which only the cognitive processing specific to the requirements of each task is occurring. Conclusions or hypotheses about the contribution of a brain area in a given cognitive process have been based on those analyses for the past few decades^46,47^. Even though most neurophysiologists are aware that correlation does not imply causation and are careful with their interpretation, numerous models of the neural basis of cognition are based on such correlational evidence^6,39^. Here, our results suggest that such correlations between neural activity and task variables are not reliable indicators of function because of the prevalence of epiphenomenal activity, and thus should not be relied upon in isolation. We believe this is especially problematic for higher associative areas of the human and nonhuman primate brain where functional roles are still unclear and undergoing intense investigation.

The present results add to emerging evidence that epiphenomenal neural activity is present across multiple brain areas^9^ and species^10^. The presence of these neural responses is puzzling. One potential explanation is that neurons are receiving inputs the area needs to perform its specialized computation (e.g. visual signals) even when the area is not involved in current task demands. Another hypothesis is that these signals are feedback from other connected areas where the critical processing unfolds. In any case, if we are to understand how brain areas collectively process information and generate complex behavior, we need to disentangle neural correlations representing a functional role from correlations representing input or feedback from other areas. To do so, causal evidence establishing a framework within which these brain-behavior correlations can be interpreted is essential.

## Acknowledgments

We acknowledge the help of Ron W. DiTullio in discussing analytical approaches and the following laboratories --Emmanuel Procyk, Konrad Kording, Yale Cohen, and Josh Gold--for insightful discussions about this work.

## Author contributions

Conceptualization: S.T., C.T., M.P. Data acquisition: S.T., C.T., J.I. Data analyses: S.T., C.T. Funding acquisition: S.T., M.P. Writing: S.T., C.T., J.I., M.P.

## Data and materials availability

The datasets generated and analyzed during the current study are available from the corresponding author upon reasonable request.

## Supplementary Materials and Methods

### Subjects

Two adult male monkeys (cynomolgus monkeys, monkey K: 7 kg, monkey L: 7 kg) participated in the experiments. All procedures complied with the Canadian Council of Animal Care and the Montreal Neurological Institute animal care committee. Over the course of a testing session (once a day), the monkeys would receive their daily amount of fluids consisting of water-diluted fruit juice for correctly performing the task. In addition, the monkeys would receive daily fresh fruits and vegetables at the end of each recording session. Each session lasted on average one hour, and no more than 1.5 hour. The physical and mental health of the monkeys was assessed daily by veterinary and laboratory staff throughout the course of the experiment. No animals were sacrificed for the purpose of this study.

### Surgical procedures

Surgical plans were prepared with the help of brain MR images obtained on a Siemens 3T scanner (TIM TRIO, Montreal Neurological Institute). T1-weighted images (MP-RAGE) with and without gadolinium enhancement (Gadovist®, Bayer, Germany) were obtained to reconstruct the 3D cortical surface and cerebral vasculature of the brains (Brainsight Vet 2.4, Rogue Research, Canada; see **Fig. 1D**). Custom Utah arrays (Blackrock Microsystems, UT) were designed on the basis of each monkey’s detailed morphology in order to cover optimally the region of interest while avoiding major blood vessels. All surgical operations were carried out under isoflurane general anesthesia and under strict sterile conditions with the help of experienced veterinary staff continuously monitoring vital signs. The head of the monkeys was positioned in a stereotaxic frame (Kopf Instruments, CA) and a midline skin incision was made to expose the dorsal aspect of the skull. The temporalis muscles were retracted ventrally and a square-shaped craniotomy (1.5cm x 1.5cm) was made in the fronto-lateral bone based on MRI coordinates using a dental drill equipped with a diamond round-cutting burr (Horico, Germany). The dura-mater was exposed and a dural flap was performed, extending ventrally. Direct visualization of cortical landmarks (i.e. the arcuate and principalis sulci) enabled identification of the pre-arcuate convexity where cytoarchitectonic area 8A of the prefrontal cortex lies in the macaque brain^48^. The array(s) were positioned over the region of interest and implanted using a pneumatic inserter held by a flexible surgical arm. The dural flap was closed with 5-0 Vycril sutures and a layer of dura regeneration matrix (Durepair, Medtronic, MN) was laid over the reconstructed flap. The bone flap was thinned with a drill and replaced over the craniotomy and secured to the skull with low-profile titanium plates and screws (DePuy Synthes, IN). The Cereport connector was secured caudally to the skull opening using eight 1.5mm diameter titanium screws with a length determined by the skull thickness as measured by pre-operative MRI. The two reference wires were inserted in between the dura and the cortex and the exposed portion of those wires and of the array wire bundle were coated with a thin layer of Quick-Sil (WPI, FL) or Geristore (Denmat, CA). The muscle, fascia and skin were closed in anatomical layers with absorbable 3-0 Vycril sutures and the Cereport connector was allowed to protrude through a small opening in the skin. The monkeys were allowed to recover for two weeks before the first recording session. No headposts were implanted in this study and no acrylic or dental cement was used in surgeries.

### Behavioral tasks

In the current study, the monkeys were trained to sit in primate chairs that did not restrict their head or arm movements. The front panel of the chair was removed allowing the monkeys to reach outside the chair with their arms. The primate chair did not impose head movement restrictions, allowing the monkeys to turn around in the chair and look in any direction (360°). The monkeys were positioned in front of a 19-inch touchscreen (ELO touch 1937l, Accutouch, CA) connected to a behavioral control computer running MonkeyLogic 2.0 (version 47, NIMH^49^) on a Windows 7 PC. The chair was positioned 7 inches from the touchscreen, such that the monkeys could reach easily at all locations on the screen. At this distance, the screen occupied 86.4 degrees of visual angle horizontally and 73.7 degrees of visual angle vertically. The two monkeys were trained on four different computerized cognitive tasks: 1) conditional association task (CAT), 2) visual discrimination task (VDT), 3) delayed match-to-sample task (DMS), and 4) spatial DMS task (DMS-s). The four tasks were performed while recording from the same neural implants over a timespan of 4 months for monkey K, and 3 months for monkey L.

### CAT

The CAT task is a computerized adaptation of the conditional association task used in Petrides et al.^17,18^. This task measures into the ability of subjects to select visual target stimuli in their environment based on instruction cues. In other words, the instruction cues require the top-down selection of particular stimuli in the environment, i.e. require the instructed allocation of attention to particular stimuli. In this earlier study, monkeys with bilateral lesions of prefrontal area 8A (the area targeted in the current study) exhibited severe impairments on this task, while sparring cognitive performance on other equally difficult tasks, such as working memory tasks. A trial was initiated by touching a white square appearing at the center of the touchscreen. Following the touch, one out of four possible visual instruction cues appeared on the screen for 1 second. After cue presentation, a delay period of random duration (0.5-1.5 sec) followed. After the delay, two targets appeared randomly at the 8 possible locations (stimuli always opposite to each other). The monkeys received a reward if they touched the correct target instructed by with the cue presented earlier in the trial. Upon selection of the correct target, the untouched target disappeared to give feedback on what target was selected, and a squirt of juice was delivered through a metal straw (Crist Instrument, MD). Before the beginning of this experiment and over the course of multiple training sessions, the monkeys had learned 36 arbitrary cue-target associations, always using the same two visual targets (a blue circle and a green star), until a threshold of 80% accuracy was reached. In any given recording session (one session per day), 4 cues were selected from the bank of 36 (2 associated with the circle, 2 associated with the star) and were interleaved in a block design, whereby a first pair of cues (1 associated with each target) was presented for half the session, and a second pair for the other half of the session. Having two cues associated with each target option permitted dissociation of selectivity for the cue vs selectivity for the association. Behavioral performance at the task was calculated using a hit rate (correct trials / (correct trials + error trials)).

### VDT

The VDT task is a computerized version of the visual discrimination task used in Petrides et al.^17,18^. This task measures a subject’s ability to select consistently a visual object in the environment regardless of contextual/instruction cues. In other words, it measures visual perceptual discrimination between stimuli. Bilateral lesions to area 8A had no effect on VDT performance. In this task, the trial structure is the same as in the CAT task (1 sec cue, 500 to 1500 msec delay). The only difference is that the cues presented were not associated with the targets. Thus, the monkeys were not instructed by the cues to select a particular visual target to perform accurately, but rather had to choose the same target on every trial, regardless of the cue presented. On every trial, a cue was randomly selected from a bank of 35 cues. This large number of cues prevented monkeys from forming false associations between cues and targets, as was previously observed with a smaller set of 4 random cues (data not shown). To ensure the monkeys selected each target (circle and star) an equal number of times, each session had a block design, whereby the first half of the trials required selection of the circle target, and the second half required selection of the star target. This order was interleaved across sessions. Monkeys quickly learned which target to select in each block through trial and error.

### DMS

The DMS task is a computerized version of the standard delayed match-to-sample task used in various lesion studies. This task measures a subject’s ability to remember after a short delay an object item in order to guide a decision after the delay. It is typically considered a “working memory” task in the neurophysiology literature^39^. The DMS task had the same trials structure as the CAT and VDT tasks. A trial was initiated by pressing a white square at the center of the screen. After the touch, one out of two visual stimuli (the circle or the star stimulus) was presented at the center of the screen for 1 sec. Following a delay period of 500 to 1500 msec, the two targets (star and circle) appeared randomly at 8 possible locations. The monkey was rewarded for selecting the target that was seen before the short delay, i.e. the star or the circle.

### DMS-s

The spatial DMS task is a spatial version of the delayed match-to-sample task. This task measures a subject’s ability to remember a spatial location in the environment in order to guide a decision after a delay. It is typically considered a “working memory” task in the neurophysiology literature. This task has the same trial structure as the CAT, VDT, and DMS tasks. First, a visual stimulus appears at one out of 4 possible locations on the screen for 1 sec. Following a delay period of 500 to 1500 msec, four targets appeared at each one of the four possible cue locations. The monkey was rewarded for selecting the target at the location that was cued earlier in the trial. To prevent the monkeys from simply leaving their hand at the cued location during the delay epoch (which would eliminate the memory component of this task), they had to place their hand back on a red dot at the center of the screen after cue offset to start the delay epoch.

### Behavioral and video monitoring

Movements explain a large portion of the neural variance in this brain area and uninstructed movements aligned with task variables can bias estimates of neural selectivity^10,38^. To account for this potential confound, we tracked head, eyes and body movements during task performance (**Fig. S2**). All behavioral monitoring devices were synchronized to the master clock of the neural recording system. All touchscreen touches by the monkeys were recorded by MonkeyLogic and marked as discrete events. In addition, we used a head-free eye tracking system to monitor eye position, pupil size, and head position (in 3D) with a 500 Hz temporal resolution (Eyelink 1000, Remote version, SR Research, Canada). This was achieved with the help a small sticker put on the monkeys’ right cheek at the beginning of each session (see **Fig. S2A**). This eye tracking signal was lost when the monkeys made very large head movements (e.g. turning 180° in the chair) bringing the sticker out of the sight of the camera. Additionally, we video-recorded the body movements of the monkeys using a video camera installed on top of the primate chair (IDS UI-3240ML-C-HQ, 30 fps, Germany; **Fig. S2A,B**).

Using video data, we measured potential biases in eyes, head or body movements that correlated with the animals’ decision on every trial (i.e. selecting the star or the circle target). We calculated the trial-averaged positions during the delay epoch, separating trials per target selected (e.g. star or circle). The distance between the average positions for star-selected and circle-selected trials is referred to as “bias” and indicate the difference in uninstructed movements made as a function of the decision taken during the delay (see **Fig. S2C,D**). Those movement biases correlated strongly with neural selectivity, persistent activity and decoding accuracy in monkey K (see **Fig. S3**) We regressed out those spatial biases from our neural metrics comparisons across tasks to control for the effect of those task-aligned movements on selectivity estimates (see details below).

### Video tracking analyses

Using the data from the video camera attached to the primate chair, we computed the motion energy for the arms and head during performance of the task. The arms and head were parsed using manually drawn regions of interests (ROI) in the Motion Energy Analysis software (MEA v4.10, F. Ramseyer 2019, https://osf.io/gkzs3/). Motion energy was computed as the number of pixels that “change” value within the ROI on a frame-by-frame basis. A “change” was defined based on a threshold crossing operation on the pixel color change to eliminate video noise (threshold = 14 a.u.).

For body movements tracking we used DeepLabCut (version 2.2)^50,51^. Specifically, we labeled 200 frames taken from 3 videos/animals (then 95% was used for training) for the following key points: right hand, left hand, eyebrow, nose, tail. The tail and left hand were visible only for monkey L and K, respectively. We used a ResNet-50-based neural network with default parameters for 150,000 training iterations. We validated with 10 number of shuffles, and found the test error was: 6 pixels, train: 307,200 pixels (image size was 640 by 480). We then used a p-cutoff of 0.8 to condition the X,Y coordinates for future analysis. This network was then used to analyze videos from similar experimental settings and identify coordinates of all key points on a frame-by-frame basis (30 frames per second).

### Neural recordings and spike detection

The neural data from the chronically implanted Utah arrays were recorded using a Cereplex Direct 96-channel neural recording system (Blackrock Microsystems, UT). The raw signal was bandpass filtered (0.3Hz to 7.5KHz) and digitized (16 bits) at 30,000 samples per second by a Cereplex E digital headstage installed on the Cereport connector of the Utah array. The digitized signal was routed from the headstage to the recording computer (PC, Windows 7) using a long, flexible micro-HDMI cable that did not impede the movements of the monkeys. For each channel, neural action potentials (or “spikes”) were detected online based on a channel-specific voltage threshold equal to 4 times the root mean square of the noise amplitude (Central Suite software, Blackrock Microsystems). The waveforms and timestamps of each spikes were saved to disk and transferred to Offline Sorter (version 2.8.8, Plexon Inc., TX) for manual sorting of spike waveforms. Well-isolated single units as well as multi-unit clusters were classified on each channel and saved for later analyses in Matlab (MathWorks, MA). On average, we recorded from 166 (SD 13.1) units (including single and multiunits) from Monkey K on each session and 134 (SD 8.5) from monkey L, or a total of 3335 units in K and 2688 units in L. We made no assumptions as to whether recorded units were the same or different ones from session to session. The number of recorded neurons was similar across tasks (**Figure 2B**).

### Single neuron selectivity

Each task taps into a dissociable cognitive process, as detailed above. The firing rate of neurons can exhibit selectivity for the cognitive representation required by each task (i.e. conditional visual selection, unconditional visual selection, working memory for object, and working memory for space for CAT, VDT, DMS and DMS-s, respectively). The selectivity of each neural unit (single or multi-unit) was assessed during the delay period using a 500msec time interval [delay start +100msec: delay start +600msec]. During this time interval, the number of action potentials was counted for each trial. Trials were separated based on whether the option 1 (e.g. star) or the option 2 (e.g. circle) was the correct answer. For the DMS-s task, there were initially four options (i.e., positions) rather than two. We therefore pooled trials into two categories: right hemifield vs left hemifield to have comparable number of trials per category with other tasks. Average firing rates over trials for each option was compared using an ANOVA with *P* < 0.01, uncorrected. Because the number of trials can influence the probability of a statistically significant result (larger N leads to lower *P* values), we randomly subsampled trials for each session to the minimum number of trials obtained in all session (87 trials per condition for monkey K, 133 for monkey L). This approach ensured that the number of neurons crossing the statistical threshold could be compared across tasks without statistical power biases.

### Common epoch across tasks: the delay epoch

Our analyses are focused on the delay epoch where no visual stimuli are on the screen and the monkeys cannot prepare a motor action (locations of targets are randomly selected during the target epoch, see **Figure 1A**). Our reasons for focusing on this task epoch are two-fold. First, the delay epoch is the most common epoch analyzed by cognitive neurophysiologists in order to dissociate sensorimotor representations from “cognitive” representations. Second, it is the only epoch that is truly comparable across the four tasks. The cue epoch has different number and types of cues across tasks, making it impossible to compare selectivity on equal grounds (CAT: 4 cues; VDT: 32 cues; DMS: 2 cues; DMS-s: 4 cues). It is also impossible to dissociate cue encoding from target encoding in a comparable manner, since the association between cues and targets is varied across tasks (1:1 in DMS, 4:2 in CAT, 32:0 in VDT).

### Correction for movement biases

As demonstrated previously, a large part of the neural variance during task performance is explained by uninstructed movements in mice^10^ and monkeys^38^. Movements that correlate with task variables inflate estimates of neural selectivity to these task variables. Since the extent of uninstructed movements varied across tasks for one of our monkeys (**Fig. S3**), to compare neural selectivity, persistent activity, auROC, and ensemble decoding accuracy consistently, we removed the neural variance explained by those movements. To do so, we ran a multilinear regression with either % tuned units, % persistent activity units, auROC, or % decoding accuracy as the dependent variable (one value per session). The independent variables were the trial-averaged uninstructed movement biases from the eyes, the cheek, the right hand, the left hand, the tail, the nose, the eyebrow, as well as the mean motion energy from the head and arms (see **Fig. S2B**). A movement bias was defined as the mean difference between the motor effector’s position (or energy) across the two target conditions during the delay epoch. The movement-controlled residuals from this multilinear regression were then used to compare selectivity, persistent activity, auROC, or decoding accuracy across tasks using ANOVAs. To obtain interpretable values on the y-axes of **Fig. 2 D-G**, we added back the intercept value (X=0) to the residuals. These final values therefore represent the estimate assuming there were no movement biases. This correction does not change results in Monkey L who does not have movement biases.

### Proportion of tuned neurons

We computed the proportion of tuned neurons for each session as: (N tuned neurons in session / N total neurons recorded in sessions). These neurons include single units and multi-units. The mean proportion of tuned neurons was compared across tasks using an ANOVA, following the same correction for uninstructed movements detailed above, applied to both monkeys.

### Persistent firing activity

Persistent firing activity was defined as statistically significant selectivity for at least 5 consecutive 100 msec time bins during the delay epoch (*P* < 0.01). Single unit selectivity was as defined above. This allowed detection of units that had a relatively persistent firing rate in favor of one of the two options during the delay. The proportion of “persistent units” was compared across tasks using an ANOVA, following the same correction for uninstructed movements detailed above.

### auROC

The area under the receiver operating curve, or auROC, is a measure of single neuron selectivity based on signal detection theory. It provides a value between 0.5 and 1, with a value of 1 being perfect binary discrimination based on the firing patterns of a single neuron. In this case, the discrimination is made between upcoming selection of the star or circle target. The function *perfcurve* from Matlab was used to compute the auROC for each neuron in a session. An average auROC was computed for each session including all recorded neurons, even the non-selective ones (which explains why the mean auROC value may appear low). The mean auROC was compared across tasks with an ANOVA, following the same correction for uninstructed movements detailed above.

### Decoding analyses

We ran decoding analyses using a machine-learning algorithm to estimate the extent of information contained in the neural firing code during the delay [delay start +100 msec : delay start +600msec]. In those analyses, we used a support vector machine (SVM) with an RBF kernel and 5-fold cross-validation. For each SVM, we only included simultaneously recorded units, hereby named “neural ensembles”, to get a more realistic estimate of information content on a msec time resolution and to account for confounding factors that can affect coding, such as noise correlations. For each cross-validation fold, non-overlapping training and testing sets of trials were defined, and the accuracy of the trained model was calculated based on the number of correct predictions on the testing set (correct predictions / all predictions). The number of trials per class (2 classes, one for each target option) was balanced using random sampling so each class had the same number of observations as the smallest class. To account for the random sampling process, we ran 30 iterations of each SVM and computed the mean decoding accuracy across those iterations. Chance performance of the decoder was obtained by randomly permuting the labels (shuffled control) before training and following the same analysis procedure. The decoding accuracy for each session was compared across tasks using an ANOVA, following the same correction for uninstructed movements detailed above.

## Supplementary Figures

**Supp. Fig. 1.**
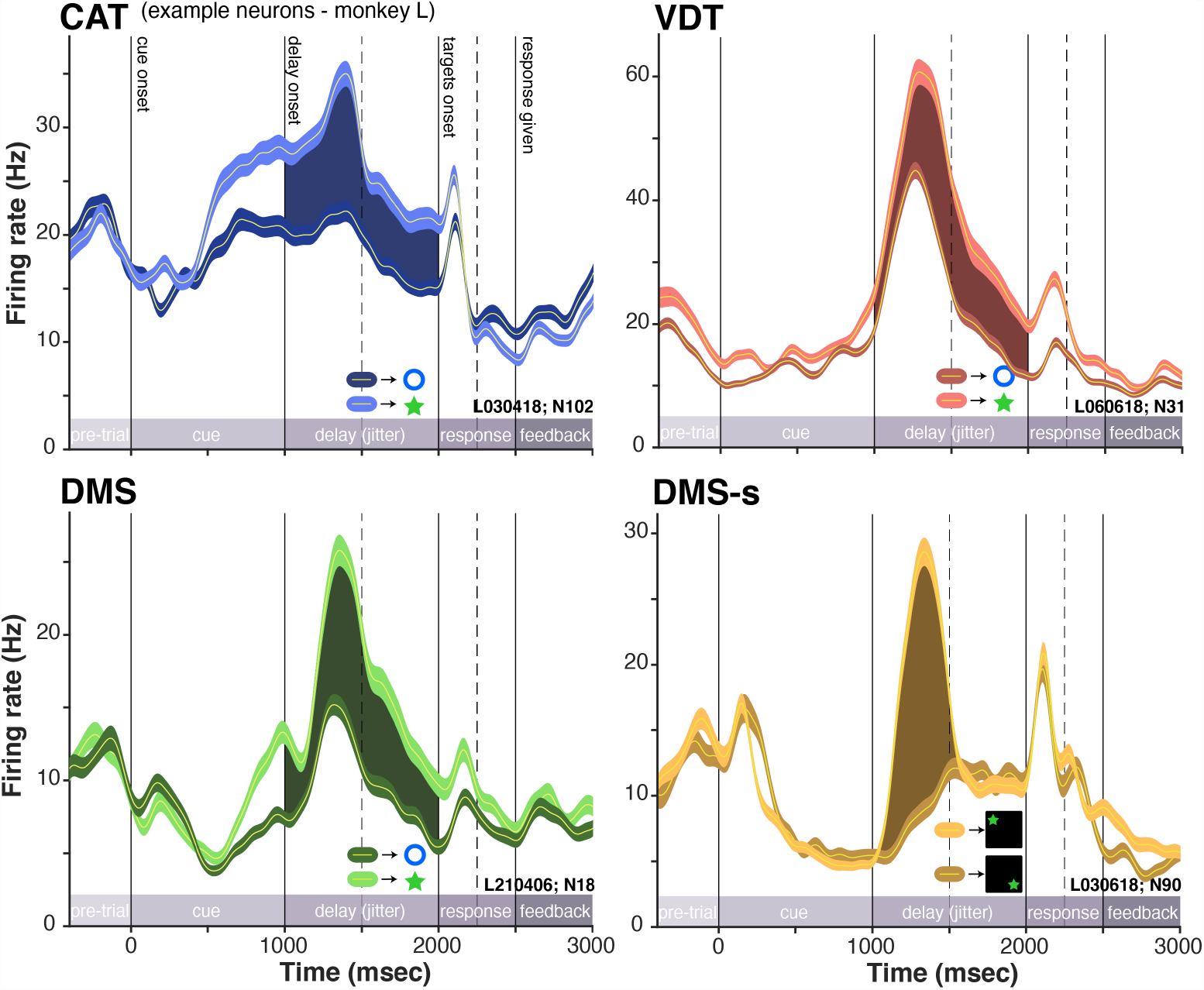
Single neuron examples for monkey L. Example single neuron spiking activity during the performance of each task, averaged over trials, for monkey L. Trials are separated based on whether target 1 or target 2 needed to be selected on this trial. This selection depended on the cognitive task. Shaded area shows significant selectivity during the delay epoch of each task, when no stimuli is on the screen. Error bars are SEM.

**Supp. Fig. 2.**
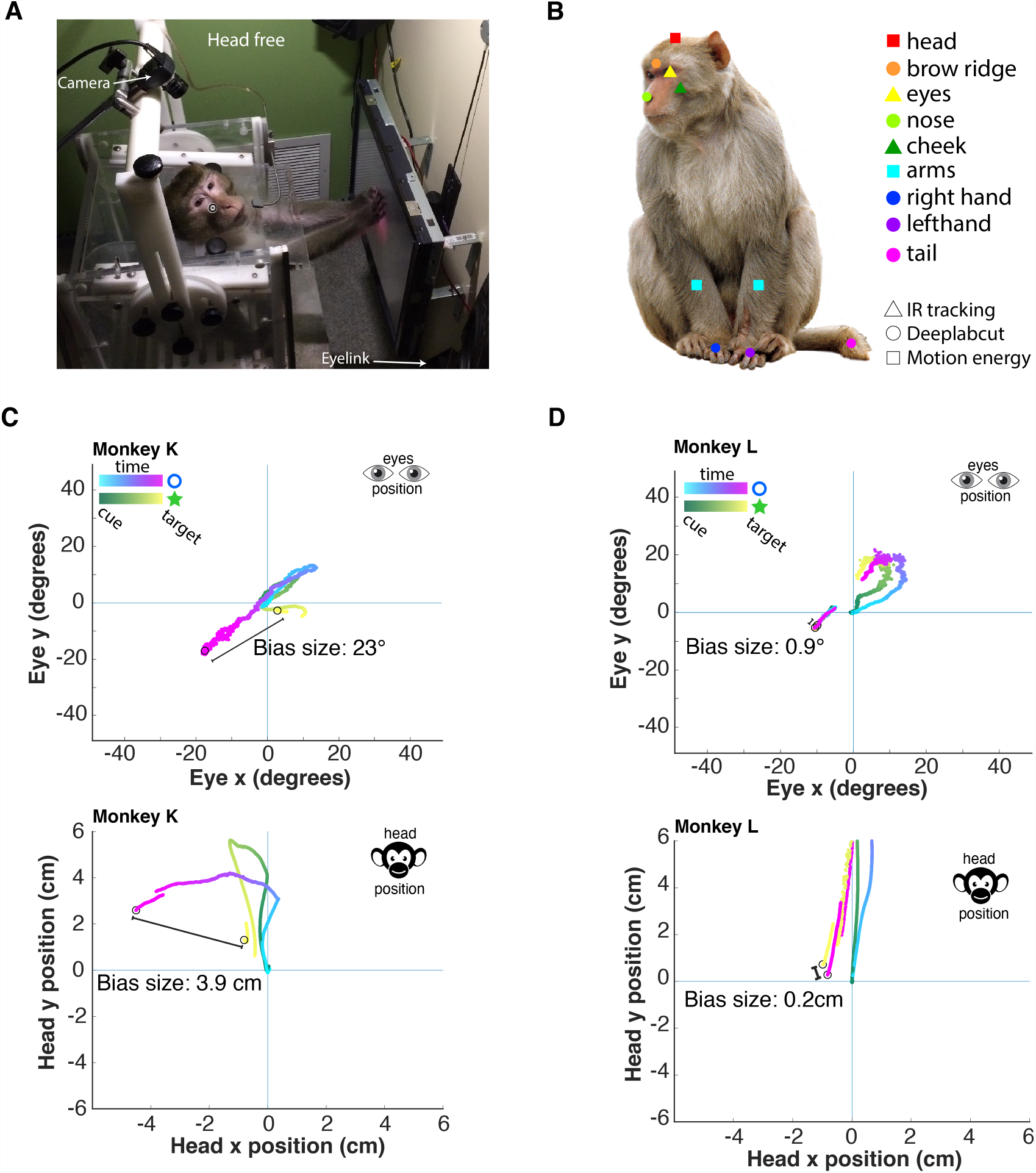
Capturing uninstructed movements. (**A**) View of the experimental setup with a head-free monkey performing a task on the touchscreen. The position of a video camera and a head-free eye-tracker are indicated. (**B**) Video tracking of macaque body movements during task performance. Various cameras and analyses techniques were used to quantify movements of 9 body parts. (**C**) Average eye (top) and head (bottom) traces during the delay of an example session in monkey K. Traces are separated based on decision to later select the star or the target. The difference between these two traces indicates uninstructed movements aligned with a critical task variable (i.e. the decision). The size of those spatial biases, calculated in degrees of visual angle for the eyes and in cm for the head, was used in the regression model to control for those movements. (**D**) Same as in (C), but for monkey L. Monkey L had almost no uninstructed movements aligned to task variables.

**Supp. Fig. 3.**
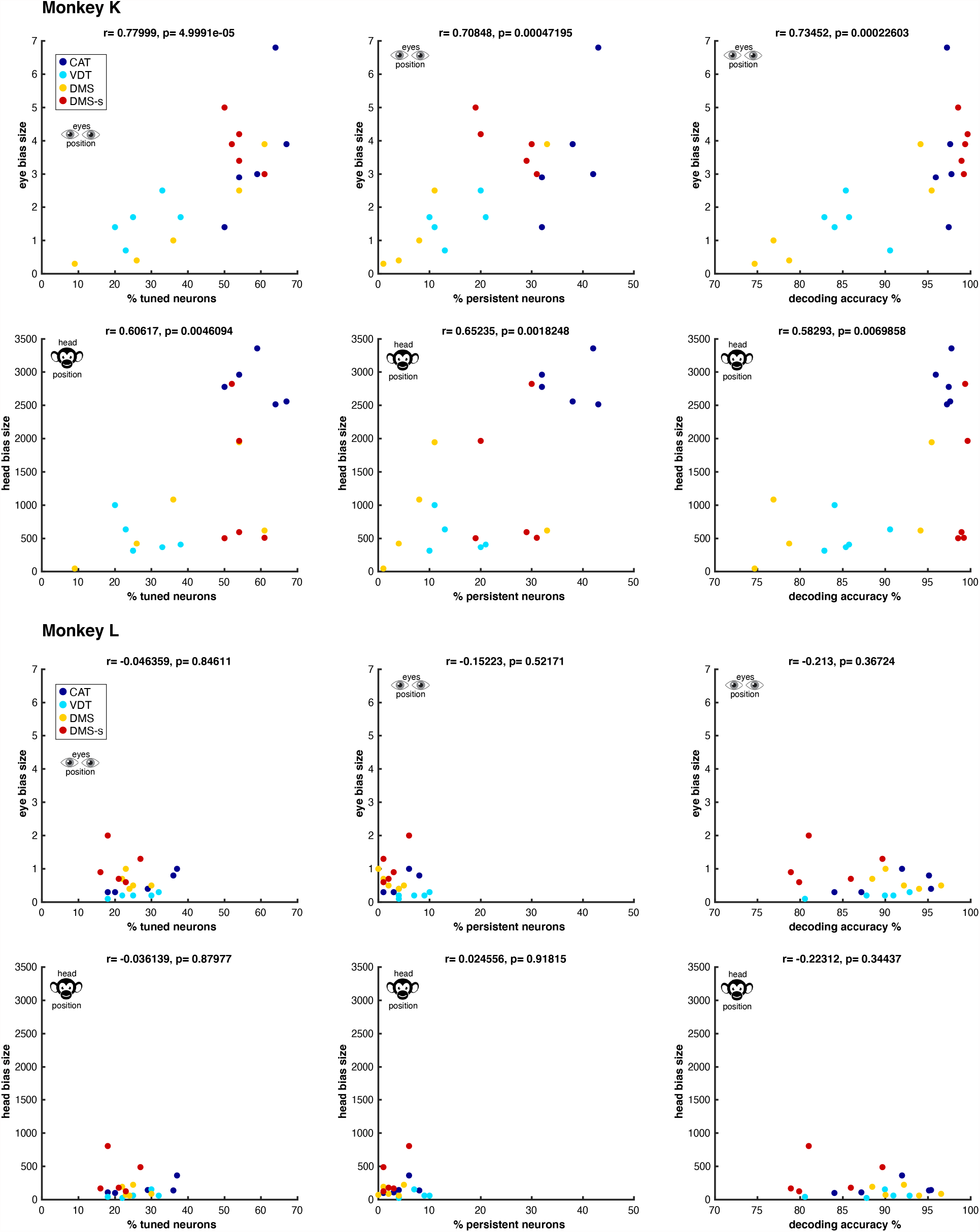
Task-aligned uninstructed movements correlating with neural metrics. Correlations between eye bias or head bias estimates, as detailed in **Supp Fig. 2**, and three neural metrics reported: % of tuned neurons (left column), % of persistent neurons (middle), and decoding accuracy (right). Note the strong biases in monkey K that vary across tasks (color legend) and correlate with neural metrics. Monkey L, in comparison, didn’t exhibit such biases or correlations with neural metrics.

**Supp. Fig. 4.**
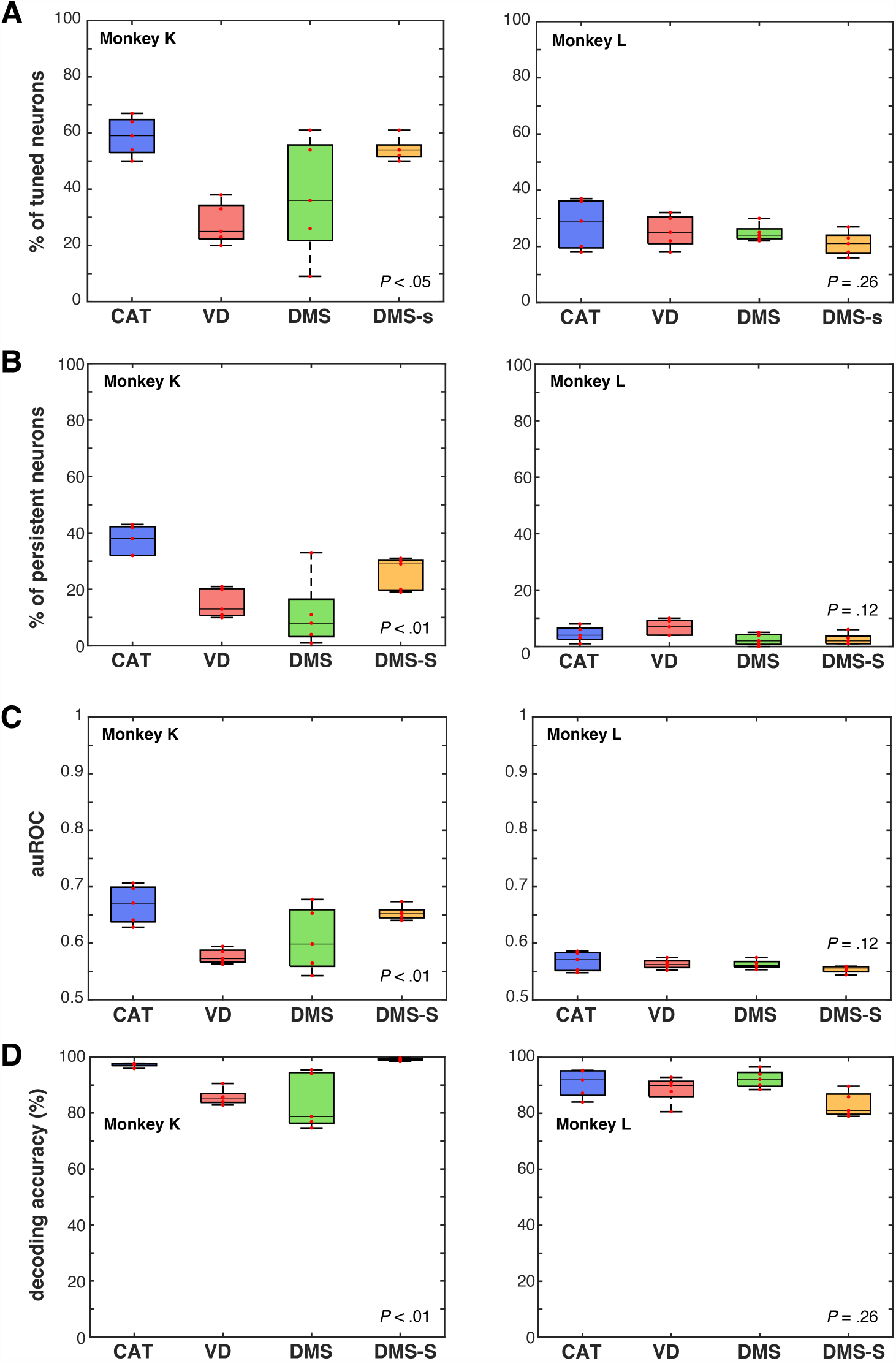
Neural metrics without correction for uninstructed movements aligned to task. (**A**) Proportion of tuned neurons when no correction is applied for uninstructed movements aligned to task variables. (**B**) Same as in A, but for proportion of persistent firing neurons during the delay. (**C**) Same as in A, but for area under the receiver operating curve (auROC). (**D**) Same as in A, but for decoding accuracy. Monkey K presented a large amount of uninstructed, task-specific movements that would inflate selectivity estimates if left uncorrected (see **Supp. Fig. 2, 3**). Conventions as in **Figure 2D-G**. Note that for monkey L who had no uninstructed movements aligned to task, the main findings hold even without movement correction.

